# Translating Brain Anatomy and Disease from Mouse to Human in Latent Gene Expression Space

**DOI:** 10.1101/2025.03.31.646307

**Authors:** Chloe Jaroszynski, Mohammed Amer, Antoine Beauchamp, Jason P. Lerch, Stamatios N. Sotiropoulos, Rogier B. Mars

## Abstract

**Background:** The mouse model is the most widely used animal model in neuroscience, yet translating findings to humans suffers from the lack of formal models comparing the mouse and the human brain. Here, we devised a novel framework using mouse and human gene expression to build a quantitative common space and apply it to models of neurodegenerative disease.

**Methods:** We trained a variational autoencoder on mouse spatial transcriptomics, and embedded mouse and human gene orthologs in the model’s latent space. We computed a latent cross-species similarity matrix for translation and compared translated maps to human ground truth evidence.

**Findings:** We established the validity of our model based on anatomical homology. Independent of species, brain areas with similar latent patterns clustered together, improving the homology of known anatomical pairs, and preserving principles of brain organisation. Importantly, brain alterations in mouse disease models predicted human patterns of brain changes in Alzheimer’s and Parkinson’s diseases. We further determined the best mouse model for the AD patients, based on how well the translations matched the patient data, across multiple models and timepoints.

**Interpretation:** Our work provides i) a quantitative bridge across evolutionary divergence between the human and the predominant preclinical species, ii) a predictive framework to help design and evaluate disease models. By highlighting which models are best suited across stages of disease, we effectively support the understanding of disease mechanisms, assist in the workflow of clinical trials, and ultimately accelerate the transformation of findings into improved human outcomes.

**Funding:** Supported by the Biotechnology and Biological Sciences Research Coundil (BBSRC) UK, the Medical Research Council (MRC) UK, the European Research Council, and the NIHR Oxford Health Biomedical Research Centre.

**Research in Context:** *Evidence before this study:* Inferences on the human brain based on findings in the mouse are backed by broad neuroanatomical similarities between the two species. However, interventions that were successful in rodents rarely translate into successful outcomes to treat human diseases. One way to address this challenge is to quantify the boundaries of translational possibilities using cross-species common space approaches. Low-level embedding spaces have been used to quantitatively compare across species, but there is currently no framework for effective translation of disease maps between species, and no tool to establish the best animal model for a given patient population.

*Added value of this study:* This study is the first to quantitatively link mouse models of neurodegenerative disease with human patient data. After validating our novel VAE-based comparative framework on anatomical homology compared to existing methods, we determine which of three Alzheimer’s Disease (AD) mouse models, and at which timepoint, best models a human patient population with mild AD. In addition, we predicted human brain changes in human Parkinson’s Disease, based on maps from mouse models of this disease.

*Implications of all the available evidence:* This approach allows one to directly evaluate mouse models based on the quality of their translation to the human. At the level of preclinical model design, the translation from human to mouse of diverse disease phenotypes will highlight the relevant targets for intervention in the mouse. This will justify the relevance of specific animal models and time courses, and therefore streamline clinical trials. By additionally translating the effect of treatment or intervention longitudinally, we can optimize human treatment plans and life-long disease management. Combined with patient population stratification, this takes an important step towards personalised translational medicine.

## Introduction

The goal of translational neuroscience is to take results obtained in the lab and use them to improve clinical practice. Today, preclinical research is producing more data, with more diversity and precision, than ever before. Genetic engineering techniques such as CrispR/Cas9 have enabled faster, more reliable manipulation of the mouse genome for creating models of human disease [1], [2], [3] and humanized mouse models have advanced in vivo investigation of human-specific responses to disease or treatment [4], [5]. Paradoxically, successful translation rates fail to match the increase in sophistication of the methods used to manipulate model genetics and compounds. The average approval likelihood in the field of neurology barely reaching the 10% mark [6], stands in contrast with the urgency of neurodegenerative conditions such as Alzheimer’s or Parkinson’s disease, recognised as the fastest rising neurological disease worldwide [7].

Surprisingly, despite the widespread dominance of the mouse model in preclinical research, a formal framework establishing the extent of the mapping between the mouse brain and the human brain is lacking. Although both brains have been mapped using a variety of methods, resulting in detailed atlases at the molecular [8], [9], [10], cellular [11], and areal levels [12], [13], neuroscientific models that allow the brain of the model species and the human to be compared in a like-for-like fashion are rare. Once anatomical mapping is achieved, translational neuroscience should seek out the quantitative translation of complex, whole brain phenotypes, across species, to provide the field with reliable predictions of effects of disease and treatment. This is a necessary and fundamental step towards improving attrition rates that undermine clinical trials.

Common space approaches have been used to describe the brains of other species in terms of a common organization feature that is comparable [14]. Connectivity fingerprints, characterizing each brain region by a unique set of connections to the rest of the brain, have bridged between primate brains using homologous white matter tracts as a common feature set. This has allowed between-species translation of cortical maps and comparisons of changes during phylogeny and ontogeny [15], [16]. The mouse brain, however, does not feature many dominant white matter bundles. Although functional connectivity with known homologous target areas has been used instead [17], a different feature with better evolutionary conservation is better suited for whole-brain mouse-human comparisons.

In this study, we present a novel approach to tackle the challenge of rodent to human translation. We use an unsupervised, generative approach in the form of a variational autoencoder (VAE), that learns to reconstruct whole brain voxel wise spatial patterns of gene expression from compressed, or latent, representations. Spatial transcriptomics have been shown to differentiate different parts of the brain [18] and datasets from both species overlap with large numbers of gene orthologs [8], [9]. Supervised region-based machine-learning approaches trained on such data have helped improve cross-species homology [19], [20], but struggle to disentangle neighbouring regions within major constituent areas of the brain. Here, after embedding in the latent space, mouse and human divergences were probed and quantitatively compared. We demonstrate that using the similarity between the latent embeddings as a basis for translation captures anatomical homologies, and highlights the conservation of hierarchical patterns of brain organisation. We show that compared to an existing supervised embedding-based method, the proposed approach provides up to 60% improvement in terms of cross-species ranked similarity decay rates. Strikingly, the approach is able to predict human patterns of brain alterations from validated mouse models of disease, in two examples of neurodegeneration, Alzheimer’s and Parkinson’s diseases. The model was further capable of determining the best match for the AD patient group, over multiple timepoints, out of three widely used mouse models of AD overexpressing the human Amyloid Precursor Protein, with different profiles of disease spread.

## Methods

### Data and preprocessing

#### Mouse in-situ hybridisation data

We used the in situ hybridisation data of an adult mouse brain from the AMBA Allen Institute database [9], aligned to the Allen Mouse Brain Common Coordinate Framework (CCFv3) [21], with a resolution of 200μm. The coronal and the sagittal datasets were downloaded and preprocessed as described in [19]. Briefly, using the pyminc toolbox in python, the sagittal and coronal images were loaded, masked, reshaped, and concatenated into voxel-wise gene expression matrices, conserving genes with multiple experiments. The matrices were log2 transformed, and the replicated experiments were averaged. Genes with more than 20% of voxels containing missing values were discarded. Below this threshold, missing values were imputed using K-nearest neighbours. Both gene sets were intersected with the set of genes common to human and mouse, resulting in a 2835 x 61315 (genes x voxels) coronal matrix, and a 2486 x 26317 (genes x voxels) sagittal matrix. Prior to further embedding or similarity computation, the data was normalized using z-score across all genes for each voxel, and de-meaned across voxels for each gene. A region-wide matrix was generated by averaging values across regions, according to a modified version of the DSURQE atlas [22] to match the AMBA hierarchical ontology [19]. White matter and ventricles were removed, and grey matter regions were aggregated to enable sufficient amounts of voxels per region for downstream training of deep learning algorithms. The resulting resolution of the atlas was of 67 grey matter regions. Annotations for 11 and 5 broad regions were also generated for visualisation purposes.

#### Resampled mouse data for independence during cross-validation

The mouse sagittal and coronal data were attributed randomly to train and test sets to maximise the independence of the data during the multiple training and validation rounds of the model for hyperparameter selection, as described in [19]. First, the coronal data was masked using the sagittal mask, for consistent spatial extent. Then, for each gene, if multiple experiments were present in the coronal set, we randomly sampled one of those for the training set, and one for the validation set. If only one experiment existed for a given gene, then we randomly assigned the sagittal or the coronal to either training or validation.

### Human microarray expression data

The human data were downloaded and pre-processed using the abagen python toolbox ***(version 0.1.3;*** https://github.com/rmarkello/abagen). To maximize spatial coverage, we generated dense gene expression maps from the full dataset provided by the Allen Human Brain Atlas (AHBA, https://human.brain-map.org; [H2012N]), consisting of regional microarray expression data from six post-mortem brains (1 female, ages 24.0–57.0, 42.50 +/-13.38). Preprocessing followed the workflow of abagen’s get_interpolated_map.

To generate interpolated maps, microarray probes were first reannotated, and probes missing a valid Entrez ID were discarded. The remaining probes were filtered based on their expression intensity relative to background noise [23]: probes below background intensity in over 50% of samples across donors were also discarded. When multiple probes indexed the expression of the same gene, we selected and used the probe with the most consistent pattern of regional variation across donors (i.e., differential stability; [24]).

The MNI coordinates of the tissue samples were updated from the original AHBA coordinates to those generated with non-linear registration (https://github.com/chrisfilo/alleninf). Samples were mirrored across hemispheres to increase spatial coverage (lr_mirror set to bidirectional). To address intersubject variation, tissue sample expression values were normalized across genes using a robust sigmoid function [25], and normalized expression values were linearly rescaled to the unit interval. Gene expression values were then normalized across tissue samples using the same procedure.

Interpolation was subsequently computed for each gene using K-nearest neighbour regression to estimate values at every voxel inside the provided mask. The resulting 1 mm resolution 3D gene expression maps with dimensions (197, 233, 189) were flattened and concatenated into a gene expression matrix spanning the 2835 common gene set, out of the total 15627 human genes. We additionally normalized the data using z-score across all genes for each voxel, and subtracting the mean across voxels for each gene.

A region-wide matrix was generated by aggregating values per region, with voxel to region mapping performed using nilearn’s NiftiLabelsMasker. For identification of brain areas, we used a modified version of the Allen Human Brain Atlas – 3D, 2020 (https://community.brain-map.org/t/allen-human-reference-atlas-3d-2020-new/405), initially based on the 2D version of the cellular resolution atlas of the adult human brain [26]. We replaced the cerebellum in this atlas by the FSL cerebellar atlas, in which bilateral regions were merged for consistency with the rest of the atlas and the mouse data, removing white matter and ventricles.

### Model architecture

We designed a model which consisted of an unsupervised variational autoencoder (VAE) combined with a latent classifier (Figure.1a). The VAE, a fully connected multi layered neural network, encoded the data onto multivariate gaussian distributions, from which the latent vectors were sampled for decoding. To avoid the issue of backpropagation bottleneck due to random sampling, the latent vector was expressed as a differentiable function: *z(x)* = *𝜇(x)* + *𝜎(x)* · 𝜖, where *x* is the input vector of the encoder, 𝜇 and 𝜎 are the output of the encoder, mean and standard deviation of the latent distribution from which we are sampling, and 𝜖 is a random variable sampled independently of 𝑥.

Simultaneously, in a supervised approach, the classifier, also a multi-layered fully connected neural network, was trained in the latent space of the variational autoencoder to recognize the 67 mouse atlas regions. The input layer of the encoder and the output layer of the decoder had a fixed number of nodes, equal to the number of genes in the study. The output layer of the classifier was equal to the number of atlas regions. The input layer of the decoder, the output layer of the encoder, and the input layer of the classifier had a number of nodes equal to the latent dimension, an adjustable hyperparameter. The number of hidden layers and the type of non-linear activation functions was also adjusted for each module. The encoder, decoder, and classifier were linked together in the loss function, which the model aimed to minimize at each iteration. The loss function consisted of two terms, to jointly train the variational autoencoder and the classifier. We used cross-entropy for the classification loss, with per-class rescaling weights to mitigate the issue of class imbalance during the training phase only:

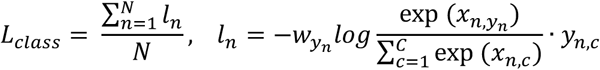

Where x is the classifier input, y is the target region index, w is the weight taken as the inverse of the frequency of the class, C is the number of classes, N spans the batch dimension.

The loss for the variational autoencoder, or Evidence Lower Bound (ELBO), consisted of a reconstruction term, to compare the output of the decoder to the input of the encoder, and a regularization term which forces the latent distributions to match the shape of the standard Gaussian distribution. We used mean squared error for the reconstruction loss and KL divergence for comparing the latent distribution to the standard Gaussian 𝒩(0, 𝐼):

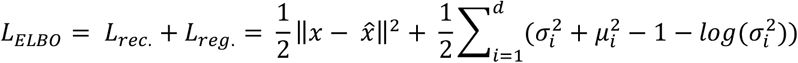

Where 𝑥 is the input vector, 𝑥̂ is the reconstructed vector, 𝑑 is the dimension of the latent space or number of latent vectors, 𝜎_𝑖_ and 𝜇_𝑖_ are the mean and standard deviations of the latent gaussian distribution over the latent space for input 𝑥. The total loss for the model was the sum of the classifier loss and the VAE loss.

### Model training and optimisation

As it was no a priori known which hyperparameters (e.g., number of layers, number of nodes) were optimal, the model was first “tuned”, meaning that training and validation was performed for different combinations of hyperparameters, using the coronal-sagittal mix described above, over 50 epochs. Then, the model with the best performance as measured in terms of accuracy in the classification task was trained and validated using the full dataset and a 80-20 split for training and validation sets, over 150 epochs.

#### Hyperparameter selection

We adjusted the type of activation function (ReLU, SiLU), the type of normalization (batch or instance), the number of hidden layers (2 or 3) and the number of nodes per layer (100, 150, or 200), the latent dimension (10,50,100,200), the amount of dropout (0, 0.1, 0.2, 0.4), the weight of the classifier loss (1, 0.5), the reconstruction weight (1, 0.5), the KL weight (0.001, 0.002, 0.005), and the maximum learning rate (10-5, 10-4, 10-3). The process was optimized using Optuna, a python based hyperparameter tuning algorithm, which sampled a subset of hyperparameters from all possible combinations, using a Bayesian optimization technique (TPEsampler), and trained the corresponding versions of the model over 50 epochs maximum with an early stopping callback to stop training after 3 epochs with no improvement. Model performance was measured using the multi-class macro-average accuracy score in the classification task.

The best model had a latent dimension of 50 nodes, an encoder with two hidden layers and 200 nodes per layer, a decoder with two hidden layers and 100 nodes per layer, a classifier with three hidden layers and 150 nodes per layer, a SiLU activation function, instance normalisation, a 10% dropout, a classifier and reconstruction weight of 1, a kl coefficient of 0.001, and a max learning rate of 10-5. We optimized the model using AdamW [27]. The maximum training accuracy over the 50 epochs was 0.865, and the maximum validation accuracy was 0.565.

### Model application

#### Data embedding and similarity

Once the optimal model was trained, we used the encoder module to project data from gene expression space to latent space, by designing the model to return the output of the encoder when calling model.predict in pytorch lightning. We transformed both the human and mouse voxel-wise and region-wise expression matrices into the latent space. Using the latent sampled vectors, we computed the similarity between the latent matrices using Pearson’s correlation, at the regional and the voxel level.

#### Comparison with previous models

We compared the performance of the cross-species comparison using our model versus using the gene expression data with no dimensionality reduction and the multi-layer perceptron developed in previous work [19].

First, we performed a 2-dimensional UMAP embedding [28] to visually highlight changes in the distribution of the data across models. Then, we computed the gene expression cross-species similarity and the MLP-embedding cross-species similarity, to highlight how the distribution of similarity values on the heatmap change with regards to the model. We quantified the differences in the regional similarity matrices using a Kolmogorov-Smirnov test.

In addition, we investigated differences in unit-scaled rank-ordered similarity profiles for each mouse region. This resulted in 67 profiles, one for each mouse region, each with a length of 129, corresponding to the number of human regions as defined by the atlas used in this study. A steeper decay rate of the ordered similarity profile highlights a more focal comparison. Differences in rank ordered profiles were tested using a two-sample Kolmogorov-Smirnov test.

#### Cross-species translation of anatomical areas

We used the voxel similarity matrices to investigate the distribution of cross-species similarity values across the whole human brain and the whole mouse brain. To optimize computation time, the voxel matrices mapped of only one human hemisphere. Therefore, each column in the matrix mapped one human hemisphere, and each row in the matrix mapped a full mouse brain. We define as translation the dot product between a vector mapping a distribution of values in one species and the similarity matrix. In the mouse to human direction, we reconstructed the brains by mirroring the data to map two hemispheres for visualisation purposes. We visualised human volumetric brain maps using FSLeyes as well as plotting tools in python (nilearn.plotting.plot_stat_map and nilearn.plotting.view_img). We provided surface-based representations using nilearn’s surface package (nilearn.surface).

First, we investigated translations using a seed-based approach, to investigate the improvement in homology of the comparison. For 35 pairs of homologous regions based on previous work and literature, we multiplied the flattened binary mask image of the ROI by the similarity matrix. This yielded a whole human brain map of values ranging from least similar to most similar to the mouse seed region. We normalised the values between 0 and 1 using min-max scaling. We first thresholded the translated maps at 0.6. Then, given that certain regions have very high, narrow profiles of similarity, we optimised this threshold by gradually increasing the value to maximize the DICE score between translated map and target region.

We quantified how well each translation was hitting its target by defining three types of scores. We defined for each target area the rate of ‘hits’ as the number of voxels above threshold in the target divided by the total number of voxels in the target. We defined the rate of ‘near hits’ as the number of voxels translating outside the target but within the same constituent part of the brain as the target area (respectively cortex, subcortex, brainstem, or cerebellum), divided by the total number of voxels in a mask of the constituent area minus the target area. We defined a rate of ‘misses’ as the number of translated voxels falling in the rest of the brain, divided by the total number of voxels in the corresponding mask. We performed an ANOVA to check for an interaction between score type and constituent part of the brain, and Connover’s posthoc test to see between which pairs of constituent brain areas the interaction was.

Then, focusing on cortical areas, we investigated gradients of cross species similarity between brain areas by computing the difference between sets of translations. To check for a primary-association gradient, we looked at the difference between the average translation of all cortical association regions (Taenia tecta, cortical amygdalar area, piriform-amygdalar area, subiculum, entorhinal area, dorsal auditory area, ventral auditory area, secondary motor area, supplemental somatosensory area, posterior parietal association areas, temporal association areas, frontal pole, and orbital area) and primary cortex areas (primary motor, primary auditory, primary somatosensory, visual areas, main olfactory bulb). We also looked at within-modality differences, i.e primary motor versus secondary motor areas, primary auditory versus dorsal and ventral auditory, primary somatosensory versus supplemental somatosensory.

To further illustrate the modality-specific effect, we computed the proportion of each primary translation falling in its corresponding target area and in each other primary cortical area, as well as in each corresponding association cortex area. For the visual cortex, the occipital lobe served as proxy for the corresponding association cortex. Similarly, we used the temporal lobe for the auditory cortex, frontal lobe for motor cortex, and parietal lobe for somatosensory cortex.

#### Cross-species translation of gene expression PCA

We performed a principal component analysis using the mouse and human ortholog gene expression data of this study, using scikit-learn’s PCA decomposition module. We masked the data to perform the PCA on the cortex only, therefore we also masked out non-cortical areas in the latent similarity matrix. We translated the first two components of the mouse, and compared them to the first two components computed directly in the human. To test for statistical significance, we transformed the maps onto the fsaverage cortical surface, and used spin tests in python using the neuromaps package [30] accounting for the spatial autocorrelation of the data. The nulls were generated in surface space using the alexander_bloch method.

#### Cross-species translation of a single Alzheimer’s disease map

We based our first disease phenotype translation on the 13-month-old APP-PS1 mouse model of Alzheimer’s disease [31]. We first coregistered the image to the template (DSURQE in CCFV3 space) used in this study. Then, we computed the cortical translation of this image, and the subcortical translation. We split the analysis to be able to run the cortical comparison on the fsaverage surface, regarded as the optimal solution in neuromaps, and the subcortical comparison in volume space.

We compared the translated map to the cortical atrophy map taken from (Lee et al, 2020), neurovault.collections:3273. We coregistered this map to the human atlas used in this study using neuromaps’ transforms module from MNI to MNI space to transform between resolutions, and symmetrised the data for consistency throughout our study. For the cortical data, we computed Pearson’s correlation and tested for significance using neuromaps.stats and the alexander_bloch method. For the subcortical data, we used Pearson’s correlation and tested for significance using the moran model for generating spatial nulls in volume space.

#### Cross-species translation of Parkinson’s disease map

To translate results from mouse to human in Parkinson’s Disease, we used the results of previous study to constitute a mask of effects in the mouse [32]. We attributed a (+1) value for regions where MPTP mice had increased tyrosine-hydroxylase expression compared to controls, and (𢈒1) value for regions where this MPTP mice had lower TH expression compared to controls. MPTP mice had reduced TH expression in basal ganglia nuclei including midbrain areas substantia nigra and substantia nigra pars compacta, caudoputamen, pallidum, subthalamic nucleus. There was an increased TH expression mainly in limbic regions including regions of the amygdala and hypothalamus. We computed the translation of this multi-region mask and compared it to the human statistical map taken from neurovault.collections:2694 [33], choosing the contrast PD>NC. We computed Pearson’s correlation and tested for significance in volume space using neuromaps.stats.compare_images and burt2018 nulls.

#### Longitudinal translation of three AD mouse models

To probe the predictive capacities of the model to determine the best match for a given patient population, we translated the three mouse models from [31], APP-PS1, hAPP-J20, and Tg2576, at respectively 5, 5 and 3 timepoints. We coregistered the mouse images to the template (DSURQE in CCFV3 space) used in this study, and masked the data for whole brain as well as cortical and subcortical translations. We multiplied the resulting vectorized maps respectively by the whole brain, cortical, and subcortical similarity matrices. For visualization only, we log-transformed the translated maps (applied log(1+x)). We used the same human AD data as previously, in MNI space, and masked consistently for whole brain, cortex, and subcortex. We computed Pearson’s correlation with each translation, and computed p-values using 1000 permutations. We plotted the profiles of correlation between the real human AD data and each translated mouse map, for the three brain masks, and across all timepoints.

## Results

### A common space based on VAE latent embeddings captures anatomical cross-species brain similarities

Using spatial gene expression data [8], [9] for 2835 genes common across the mouse and the human, we trained an unsupervised neural network to learn low dimensional representations of the mouse transcriptomics data. Specifically, we used a variational autoencoder (VAE) architecture tasked with reconstructing the gene expression data based on the latent representation of the input, and a latent classifier tasked with recognising the regions of the Allen Mouse Brain Atlas based on the latent representation of the input (Fig. 1 (a)). After training, the encoder was used to project both mouse and human spatial transcriptomic data into the latent space (Fig. 1 (b)). This transformation enabled voxel-wise gene expression profiles – originally represented as N-dimensional vectors – to be compressed into M-dimensional latent vectors (M<<N) for both species. By establishing this common transcriptomic space, the M-dimensional latent representations facilitated direct comparisons and cross-species translations.

**Fig. 1.**
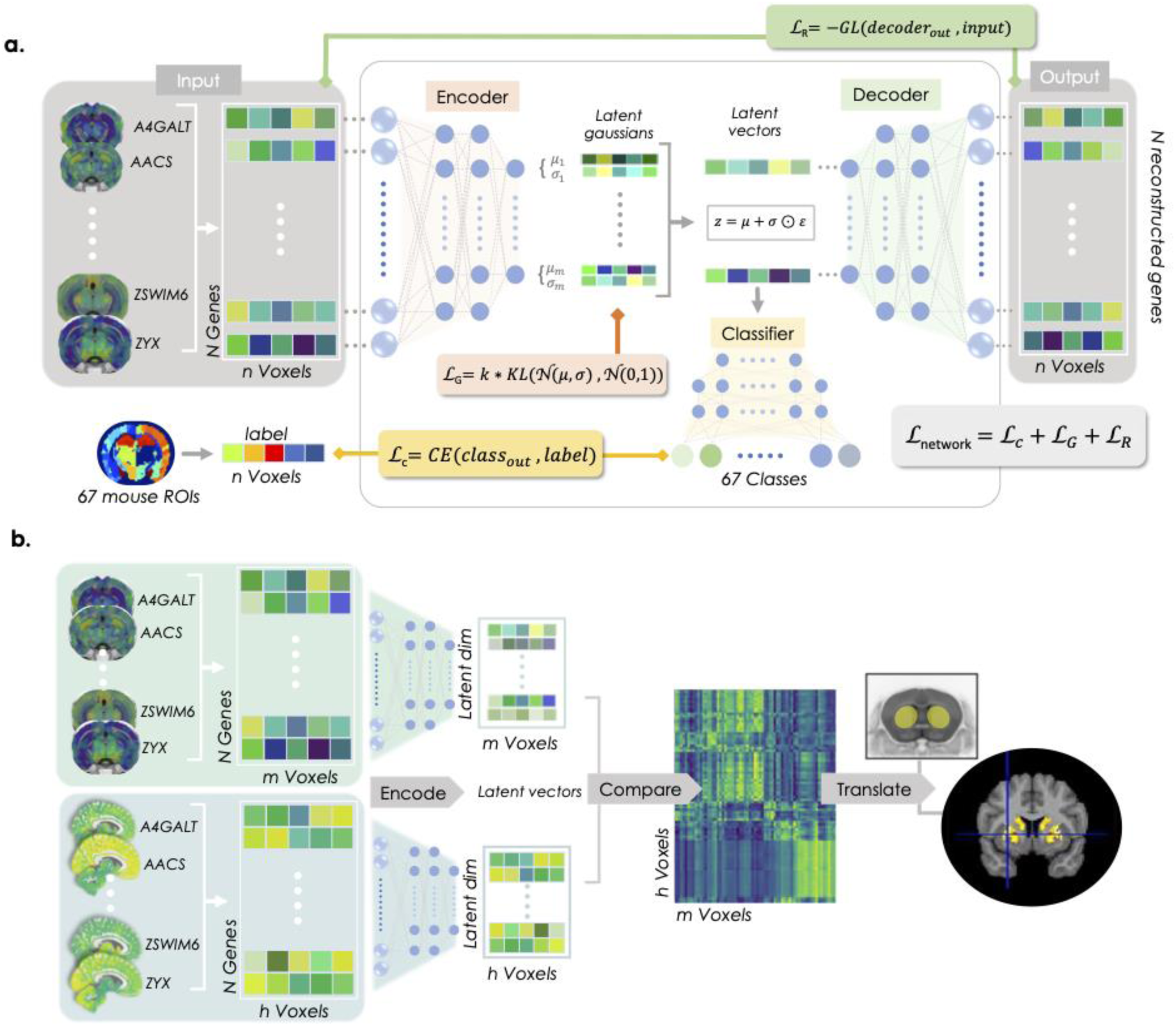
A Variational Autoencoder trained on mouse gene expression data leads to quantitative translations between the mouse and the human. **a** – Tripartite model architecture with encoder, decoder and latent classifier trained on the mouse data using N=2835 ortholog genes; GL: Gaussian likelihood; KL: Kullblack-Leibler divergence; CE: cross-entropy. **b** – Application: the encoder is used to embed the mouse and human data into the latent space of the model for similarity computation and cross-species translations.

### Alignment of cross-species distributions in 2D-Umap projections

To investigate the performance of the VAE in terms of cross-species alignment, we first examined its ability to preserve within-species regional organisation, while bringing the two species closer together for effective comparison. To this end, we project the relevant mouse and human data into two dimensional UMAP visualizations. To examine this aspect with respect to alternative methods, we evaluate three approaches – the naïve approach using the original gene expression data, the multi-layer perceptron (MLP) derived embeddings [19], and the VAE embeddings (Fig 2. (a)).

**Fig. 2.**
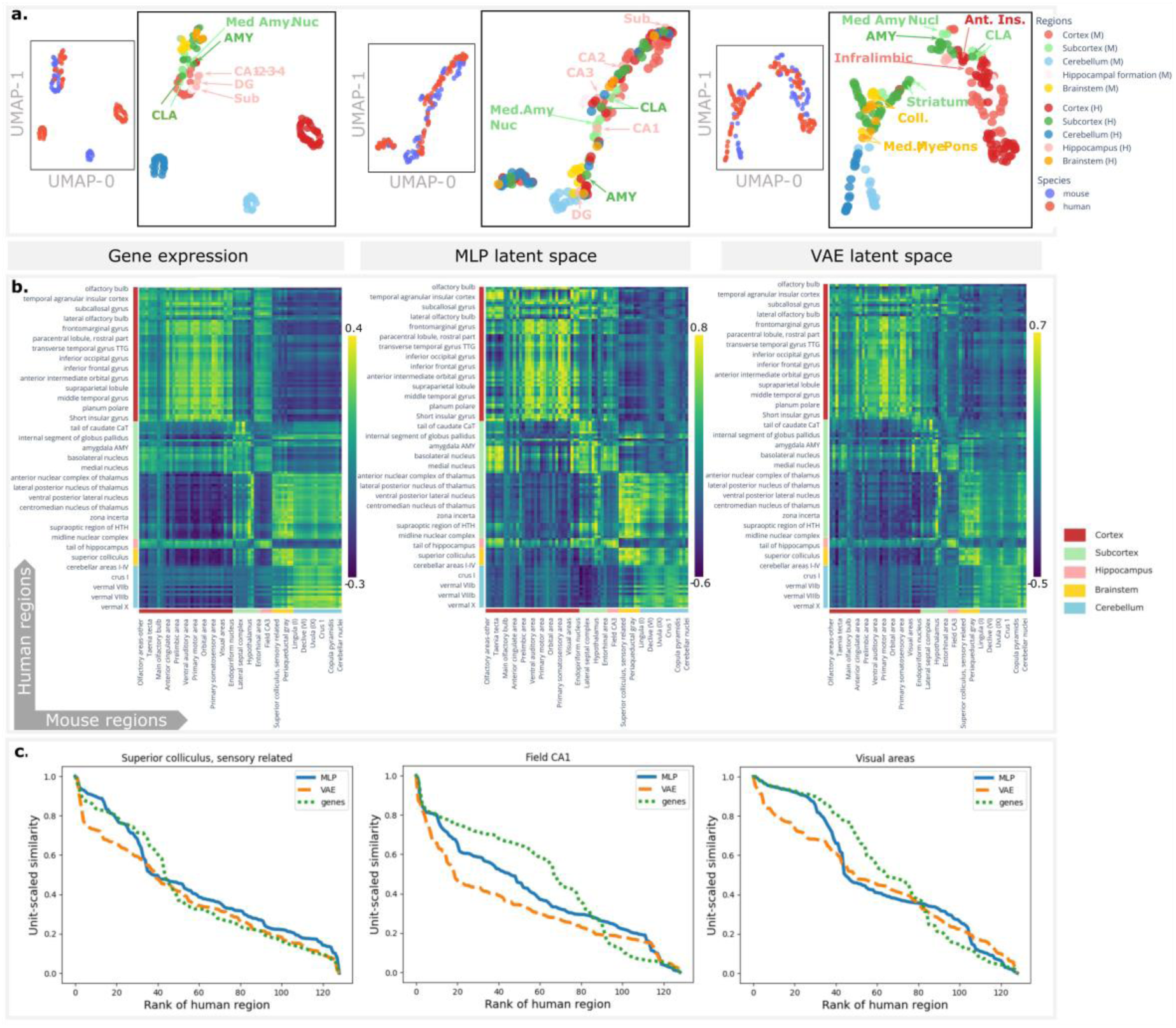
The VAE improves the cross-species comparison compared with the gene expression and the MLP approaches. **a** – 2-D Umap for visualisation of regional data distributions, where each dot represents a mouse or human ROI, as defined per the atlases described in the methods. The VAE achieves the best cross-species alignment and regional clustering within species. DG=Dentate gyrus, Sub=subiculum, med. amyg. nucl=medial amygdalar nucleus, Cla/CL=claustrum, Med=medulla, mye=myencephalon, ant. ins = anterior insula. **b** – Heatmaps of region-wise cross-species similarity where axes are anatomically ordered according to major brain subdivisions (human and mouse regions on vertical and horizontal axes respectively). **c** – Unit-scaled, rank ordered similarity profiles for 3 regions and for each method. The VAE displays a steeper slope of the curve at the origin, and a lower rank at 0.75 similarity, highlighting the improvement of the VAE in terms of narrowing the similarity profile for these regions.

In the original gene space (Fig 2 (a) - left), alignment is weak. Clustering is dominated by species rather than anatomical correspondence: while some human regions (e.g. hippocampal formation, claustrum, amygdala) align with their mouse homologs, most cortical, subcortical, and cerebellar areas cluster strictly by species. The MLP representation (Fig. 2 (a) - middle) reduces inter-species distance but at the cost of within-species alignment. Although some homologous structures (e.g. Claustrum) remain in close proximity, overall organisation is degraded. A number of human brain regions are scattered across unrelated domains, such as the hippocampal subfields which intermingle with cerebellar and cortical regions. This loss of structure may result from the sensitivity of the MLP to mouse-only training prior to cross-species embedding.

In contrast, the VAE latent space (Fig 2 (a) - right) achieves simultaneous within-species clustering and cross-species alignment. In particular, the hippocampal subfields cluster closely within and across species and are surrounded by human and mouse limbic structures; brainstem and subcortical areas are well aligned across, as well as within species. Some cortical and cerebellar areas still cluster primarily by species: this is consistent with expected divergences between phylogenetically distant species, but warrants further investigation.

This demonstrates that the VAE latent space embedding presents an improvement in cross-species alignment that, importantly, preserves anatomical segregation.

### Quantitative comparison of regional alignment

To further investigate the improvement of regional alignment of areas across mouse and human brains using the VAE, we computed for each approach the cross-species regional similarity matrices using Pearson correlation (Fig 2 (b)). We tested the improvement using a Kolmogorov-Smirnov test on the distribution of the similarity values across approaches and additionally investigated the differences in terms of rank ordered similarity profiles of each region (Fig 2 (c)).

The heatmaps highlight the progression of the cross-species comparison between the three approaches. On one hand, while the large high similarity clusters in the gene similarity matrix (Fig 2 (b) – left) reflect the major anatomical subdivisions, they do not identify specific cross-species homologs. Mouse cortical regions are mostly similar to human cortical regions but have high similarity to the hippocampus and to subcortical areas; the mouse subcortex is mostly similar to the human subcortex but also to areas of the brainstem; the mouse cerebellum is mostly similar to the human cerebellum, brainstem and subcortex.

The clusters of high similarity in the MLP similarity matrix are less uniform than the gene similarity matrix, reducing the number of cross-species matches for any given region. However, matches still occur across major anatomical subdivisions, between cortex and subcortex, brainstem and subcortex, cortex and hippocampus, or cerebellum and subcortex. This is consistent with the poorer alignment observed in the MLP UMAP representations in figure 2a.

In contrast to the gene and MLP matrices, the VAE similarity matrix is both more focal and anatomically accurate. Mouse cortical regions no longer have high similarity to regions outside the human cortex, mouse subcortical similarity also remains within the human subcortical areas, and the same can be seen for hippocampal fields and cerebellum. Additionally, the number of high similarity correspondences for potential cross-species homologs is further reduced.

On a general standpoint, the Kolmogorov-Smirnov test confirmed that the distributions of similarity values in the matrices were significantly different (p=5.77e-6). More specifically, the rank ordered, unit-scaled similarity profiles highlight the level of improvement for each region. A steep decay rate indicates a focal cross-species similarity, while a flat decay profile indicates that the region is highly similar to many regions in the other species. Figure 2 (c) illustrates this in 3 regions where the VAE shows steeper decay rates compared to the gene expression and MLP approaches. The difference between the ranks of each profile at a 0.75 similarity level was on average in favour of the VAE by 2.7 percent or over 3 regions. Comparing the improvement of the MLP versus genes (4.63%) and the VAE versus genes (7.38%), on average the VAE showed a 60% improvement compared to the MLP.

Taken together, these observations demonstrate the efficacy of the VAE in representing regional data from both species using a common, low-dimensional latent space, with a balance between cross-species alignment and within-species clustering of brain regions.

### Regional translations: retrieving known homologs and exploring new correspondences

#### Translation between known homologs

Having established the validity and specificity of the VAE latent space, we used it to compare the mouse and human brain. As a starting point, we investigated the translatability of specific regions of the mouse brain.

We first investigated the cross-species translation of the hippocampus, caudoputamen, thalamus and hypothalamus. These areas are known to be close in anatomical space, i.e., they are located closely together in the brain but are thought to be far apart in terms of gene expression [8]. Figure 3 (a) presents a translation of each of these regions, in which a binary mouse ROI, defined using the Allen Mouse Brain Atlas, was translated to human (mouse ROI multiplied by the mouse-human similarity matrix to create human brain similarity map of the mouse region, scaled to [0 1] range and thresholded as described in methods). All regions match best to their human homologs with high selectivity, demonstrating that these regions translate well using the VAE latent space.

**Fig. 3.**
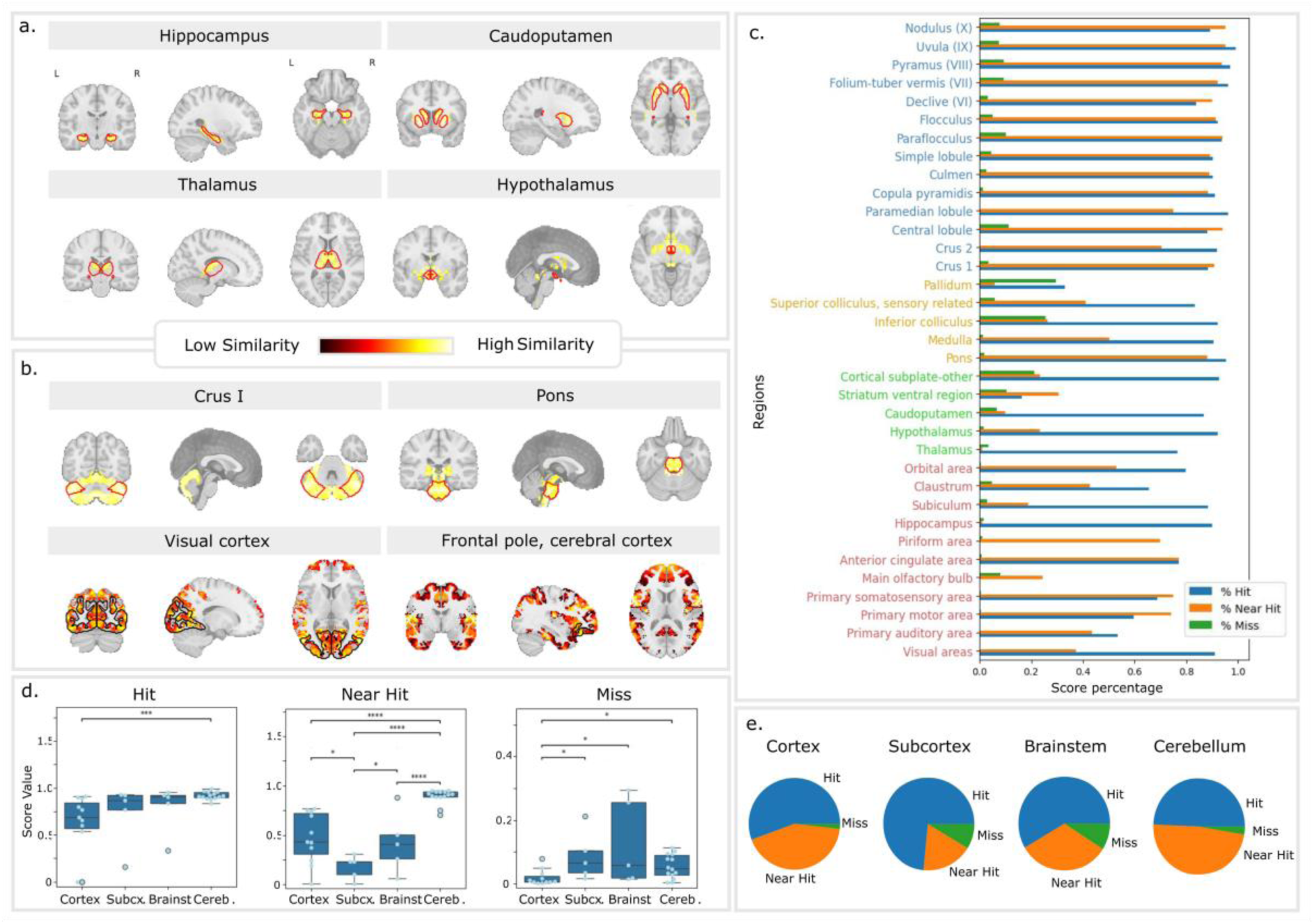
Cross-species translation between homologs. **a** – thresholded maps of translations from hippocampus, caudoputamen, thalamus, and hypothalamus. Left and Right hemispheres are indicated on the first plot and are the same for all plots. The red line indicates the target human area. **b** – thresholded maps of translations from crus 1, pons, visual cortex, and frontal pole, with the human target indicated by a continuous red or black line. **c** – proportions of the target region containing voxels with high similarity in the translation of the mouse homolog (% hit), proportion of the larger anatomical structure containing high similarity voxels (% near hit), proportion of the rest of the brain containing high similarity voxels (% miss), for 35 homologous regions. **d** – significant differences between hit, near hit, and miss rates in the different constituent parts, according to connover’s post-hoc test. **e** – proportion of score types in each constituent part.

We next investigated a wider range of areas across all major subdivisions of the brain. Figure 3b shows the translations of mouse Crus I (a cerebellar area), the pons (a brainstem area), the visual cortex (a primary cortical area), and the frontal pole (an association area of the cortex). All areas translate well in terms of having high areas of similarity in the homologous part of the human brain. However, in some cases, high similarity is also found in areas that are outside the direct homolog. For areas where this is the case, additional voxels with high similarity scores tend to be found in the same constituent parts of the brain (e.g., cortex, subcortex, cerebellum, brainstem). This is particularly the case for the neocortical areas. Both for the visual cortex and for the frontopolar area, voxels with high similarity are found outside the target area but still inside the neocortex.

To quantify this aspect, we used predefined pairs of mouse-human homologous areas based on the literature. We also defined anatomical constituents for each brain region based on the anatomical area it belonged to (cortex, subcortex, brainstem, cerebellum). We then calculated for each mouse area the percentage of voxels within the homologous human area that had a high similarity (“hit”), the percentage of voxels within the rest of the constituent part that had a high similarity (“near hit”), and the percentage of voxels in the rest of the brain that showed a high similarity (“miss”). Figure 3 (c) shows these three scores for the 35 homologous areas. Across the whole brain, the rate of voxels falling beyond the major anatomical constituent is low. However, the pattern of scores differs across the different constituent parts of the brain. The more focal translations are found mainly in the cortex, subcortex, and brainstem, potentially highlighting a better homology between species in those areas. In 14 regions across these constituent parts, there is a higher rate of hits than near hits, and hardly any misses, which models how the similarity decreases as the distance to the target increases. In the cerebellum, the pattern is rather that there are similar proportions of hits and near hits, meaning that the translations may not be discriminating between the cerebellar subregions as well as for other constituent brain areas. An ANOVA on hit, near hit, and miss percentages for each brain area with factors SCORE TYPE (hit, near hit, miss) and CONSTITUENT PART (cerebellum, brainstem, subcortex, cortex) shows a significant interaction (*F*(6,93)=10.85, *p*<0.001). Connover’s posthoc test highlights the significant pairwise differences for each score type (Fig. 3 (d)), while the pie-plots in Fig 3 (e) highlight the relative proportion of each score type, each constituent part taken independently.

The fact that the subcortex and brainstem highlight the most focal mouse-human translations is consistent with the idea that these areas are conserved in the mammalian lineage. One could have expected the same in the cerebellum, but the score pattern in the cerebellum does not necessarily mean that the translations are poor in this area. Perhaps the cerebellum is overall more different between species than the brainstem and subcortex, and would benefit from a dedicated analysis. Another reason could be that the anatomical atlas used as first intention in this study may not be the best suited for cross-species translations, and that the similarities could lie within functional boundaries (such as defined by [34]) instead.

Similarly, the variability in hit and near hit scores in the cortex should not necessarily be taken to mean that translation is worse in this part of the brain. The neocortex has expanded dramatically in the primate lineage and contains many more distinct areas in the human than in the mouse. While the pattern of organisation can be similar, the expansion that took place in the human brain results in subsequent diversification of areas [35]. Cortical areas resulting from brain expansion do not necessarily have corresponding diversity in their patterns of gene expression, meaning that they could be classified as similar to mouse areas, even if they have no direct homologs in the mouse. To further investigate this issue, we focused on the translation of cortical areas in more detail.

To further investigate this issue, we focused on the translation of cortical areas in more detail.

#### Neocortical translations

First, we investigated the translation of modality-specific primary areas. We showed previously that the visual areas translate accurately across species, with the highest percentage of hit voxels among primary cortices (Fig. 3 (c)), while primary motor, primary somatosensory, and primary auditory areas displayed a much higher rate of off-target voxels (near hits). There is, however, some structure to the location of these translated voxels. To illustrate this, Fig. 4 (a-c) shows the annotated translations of these areas on the human brain, together with the outline of the Allen Human Brain Atlas regions [29].

**Fig. 4.**
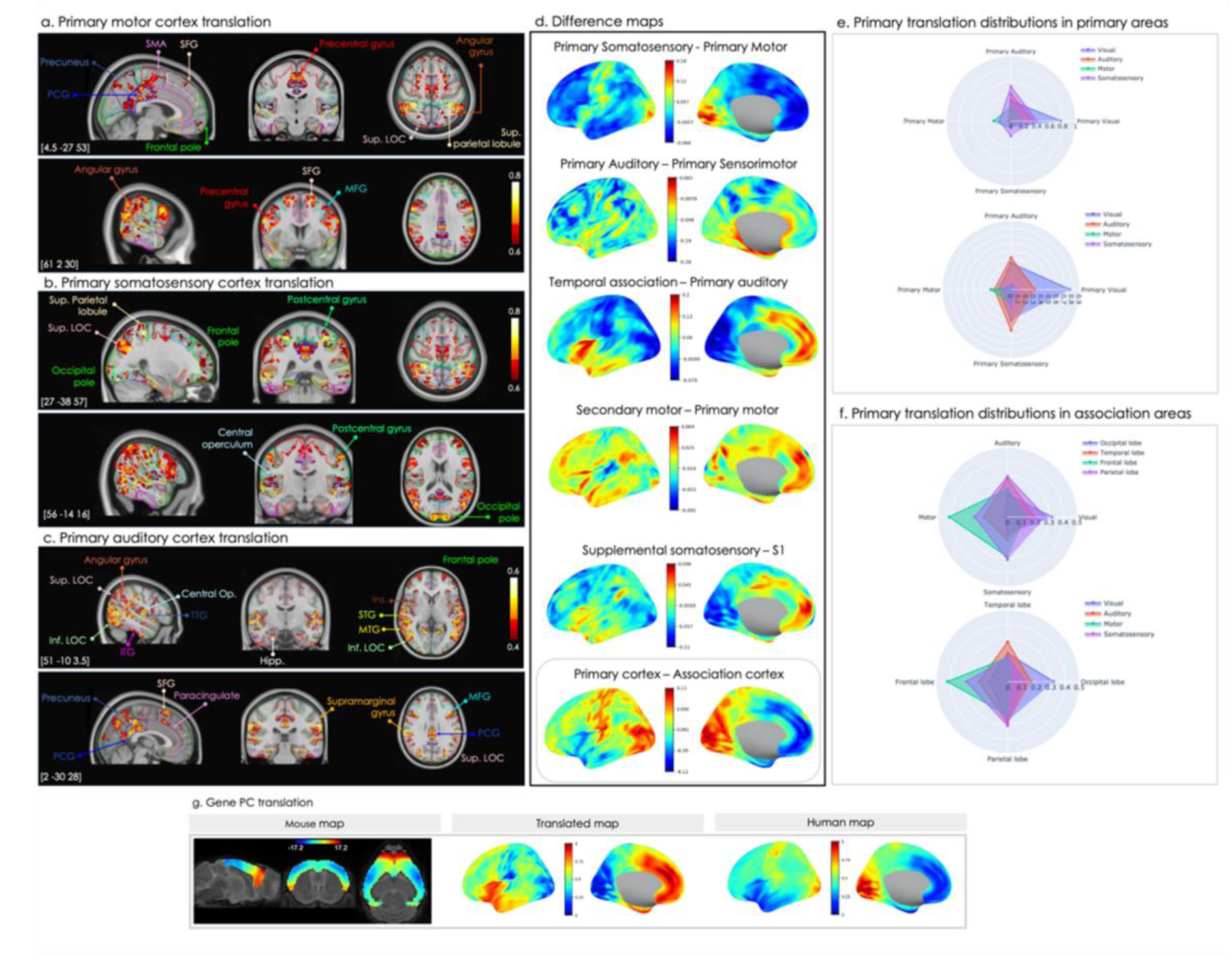
Cross-species translations in neocortical regions. a, b, **c** – Thresholded voxel-wise volumetric translations from mouse primary motor, primary somatosensory, and auditory cortex. **d** – Surface representation of difference maps highlighting patterns of similarity within and across sensory modalities. **e** – Average distribution across human primary areas of translated voxels from mouse primary areas. The top row can read as ‘Where does this mouse region land in the human brain?’ and the second row can read as ‘From which mouse area does this human area receive voxels?’. **f** – Average distribution across frontal, temporal, occipital and parietal lobes (association cortices) of the mouse primary translations. **g** – Translation of the gene expression PCA of the mouse and comparison with the gene expression PCA computed in the human.

In the translation from the mouse primary motor area, high similarity in the human precentral gyrus is found mainly in two areas. The area on the medial surface of the cerebrum (Fig. 4 (a), top panel) is responsible for foot and toe movement, while the area adjacent to the sylvian fissure (Fig. 4 (a), bottom panel) is responsible for face, eye, lip and tongue movement. The mouse primary motor area also displays high similarity with the posterior cingulate, precuneus, and superior parietal lobule, with the lateral occipital gyrus and the angular gyrus, and with the frontal pole, middle and superior frontal gyrus, and the posterior temporal gyrus.

The translation of the mouse primary somatosensory area broadly overlaps with the translation of the primary motor area, however with a cluster of similarity of larger spatial extent in the postcentral gyrus adjacent to the superior parietal lobule (Fig. 4 (b), top panel), possibly involved in the sensation of the hand and fingers, and in an area of the postcentral gyrus adjacent to the supramarginal gyrus (Fig. 4 (b), bottom panel), involved in sensations of the face, mouth, or nose. Outside the postcentral gyrus, clusters found in the somatosensory translation that do not overlap with the motor translation are located in the central operculum (as defined in the Harvard-Oxford Cortical Structural atlas – Fig. 4 (b), bottom panel) and the parietal operculum, involved in somatosensory processing of laryngeal areas, as well as in the occipital pole.

The primary auditory cortex translation (Fig. 4 (c)) intersects with the somatosensory and motor translations in the posterior cingulate, the lateral occipital cortex, the occipital pole, middle frontal gyrus, and central operculum. It differs in the frontal pole, precuneus, angular gyrus, supramarginal gyrus, parietal operculum, superior frontal gyrus, and temporo-occipital middle temporal gyrus. Importantly, high similarity specific to the auditory translation is found in areas involved in primary and higher order auditory processing: the temporal pole, the planum polare, the anterior superior, middle and inferior temporal gyri, as well as in Heschl’s gyrus, in the lingual gyrus, the temporal fusiform gyrus, the parahippocampal gyrus, and the posterior insula.

By looking at the differences between these primary translations, we highlight further patterns of similarity (Fig. 4 (d), rows 1-3). We observe that the primary somatosensory area shows greater similarity to primary and insular areas than the primary motor area, which is comparatively more similar to frontal, temporal, and parietal association areas (Fig. 4 (d), row 1). The auditory translation has a higher similarity to temporo-insular areas, areas typically involved in higher order auditory processing compared with the somatosensory translation (Fig. 4 (d), row 2), which highlights a parieto-frontal pattern of similarity, including areas more typically involved in higher order somatosensory processing.

Looking within modalities (Fig. 4 (d), rows 3, 4, 5), we can see that the auditory primary-association difference map (computed as the difference between temporal association and primary auditory cortex translations) is directed broadly from non-specific primary areas to anterior temporal, insular and medial frontal areas. The motor difference map is more diffuse across the cortex, from more focal primary areas – with emphasis on auditory – to parietal, fronto-temporal, and medial frontal areas. The somatosensory difference map, conversely, shows an emphasis on visual primary, and is directed more focally towards fronto-temporal and parietal areas.

On average (Fig. 4 (d), bottom row), we highlight a primary-association pattern of cross-species similarity, by creating a difference map between the average translation of mouse primary regions (motor, somatosensory, visual, auditory) and the average translation of mouse association regions to the human. On average, the mouse primary areas have higher similarity to human primary areas (in red in Fig. 4 (d), bottom row) than to human association areas, and mouse association areas have higher similarity to human association cortices (in blue in Figure 4 (d), bottom row) than human primary cortices. This is in line with previous results showing that a substantial portion of the variance in gene expression among cortical areas can be described as a gradient between primary areas and association cortex (Burt et al., 2018).

To quantify the structure underlying these patterns of cortical similarity with respect to modality, we looked at the distribution of translated voxels from each primary modality within major anatomical areas (Fig. 4 (e, f)). First, looking at primary areas only (Fig. 4 (e)), we investigated the proportion of voxels within each primary area in the human cortex originating from each translation (top figure), and alternatively at the proportion of voxels in each primary translation landing within each primary region in the human cortex (bottom figure). Both cases confirm the strength of the visual translation over other modalities, and similar patterns for the motor translation with smaller but consistent rates of voxels translating to target. The auditory and somatosensory translations show a degree of trade-off, depending on the angle of discussion. On one hand, we highlight that while the primary area receives more voxels from the somatosensory translation than from the auditory translation, the primary somatosensory area receives voxels mainly from the somatosensory translation. On the other hand, we highlight that the translation of the mouse auditory region goes first to auditory regions, and that the translation of the somatosensory area sends more voxels to the auditory and visual regions than to the somatosensory areas.

Looking at the distribution of primary translated voxels with respect to association areas, we highlight that primary translations predominantly remain within their corresponding association areas (Fig. 4 (f)). This is particularly clear for the visual and motor areas. The parietal lobe appears more multimodal, but still retains most voxels from the somatosensory modality. The auditory modality projects to both the parietal lobe and the temporal lobe, but we can see that the temporal lobe receives primarily voxels from the auditory regions. This confirms that there exists a within-modality pattern of cross-species similarity.

Cortical areas thus seem to translate from the mouse to the human in a reliable way. The fact that the human cortex is larger, more complex and subdivided could explain why single mouse regions tend to show high similarity to a series of human regions. This one-to-many mapping shows two overlapping principles of organization. First is a translation gradient between primary and association cortices (Fig. 4 (d) – bottom row); mouse primary areas are more similar to human primary areas, while mouse association areas are more similar to human association areas. Second is a within-modality gradient, whereby primary areas tend to show high similarity to areas of association cortex generally associated with processing the same type of information, as demonstrated in Fig. 3 (f) and illustrated in Fig. 4 (d) (rows 4 and 5).

#### Translating a cortex-wide scalar map

Moving beyond regional translations, we performed a first cortex-wide translation of a quantitative map (Fig 4 (g)). Using the mouse-human similarity matrix, a scalar map in the mouse cortex was projected onto the human cortex by multiplying with the similarity matrix.

In humans, the first principal component of gene expression maps across brain-specific genes has been shown to correlate with the primary-association gradient of cortical hierarchy (Burt et al, 2018). We tested whether this was mirrored in the mouse brain, and if the translation of the first principal component of the mouse gene expression data matched the human gene expression PC1.

We ran the principal component analysis in both species on the subset of mouse-human ortholog genes used in our analysis. The second component highlighted a pattern consistent with a primary-association gradient, replicating findings from [36]. For the human PCA, the first component was most consistent with a primary-association gradient and correlated significantly with the gene expression PC available from the *abagen* toolbox [37] (r=0.70, p<0.001). We thus performed the translation of the mouse second component, and since this translation was restricted to cortical areas only, we multiplied the mouse data by the cortical part of the latent similarity matrix. We compared the translated map with the human gene PC1, using neuromaps’ statistical tools for spin tests accounting for spatial autocorrelation. The translated map correlated significantly with the human map (r=0.9, p<0.001, irrespective of arbitrary sign of the PCA).

Having achieved a formal anatomical comparison of the mouse and human brain on the basis of the similarity of their embeddings in VAE latent space, we now move on to our second goal: translating disease phenotypes between species.

### Translation of disease phenotypes across species

#### Translating a map of Alzheimer’s disease

To demonstrate the use of the latent space to translate disease phenotypes, we performed a cortical translation of the mouse AAP-PSI model at 13 months, corresponding to a fairly advanced stage of the disease [31] (Fig 5 (a)). This model overexpresses mutant forms of the human amyloid precursor protein resulting in plaque formation due to amyloid beta deposition in the brain, first in the isocortex and with regional specificities. For comparison, we obtained a statistical map quantifying the effects of Alzheimer’s disease compared to healthy controls in the human, from a study investigating networks of regions presenting grey matter volume decline, or synchronised degeneration networks [38]. This study highlighted the posterior cingulate cortex and the hippocampus as vulnerable epicentre regions in Alzheimer’s disease, and their degeneration networks as a marker of disease progression.

**Fig. 5.**
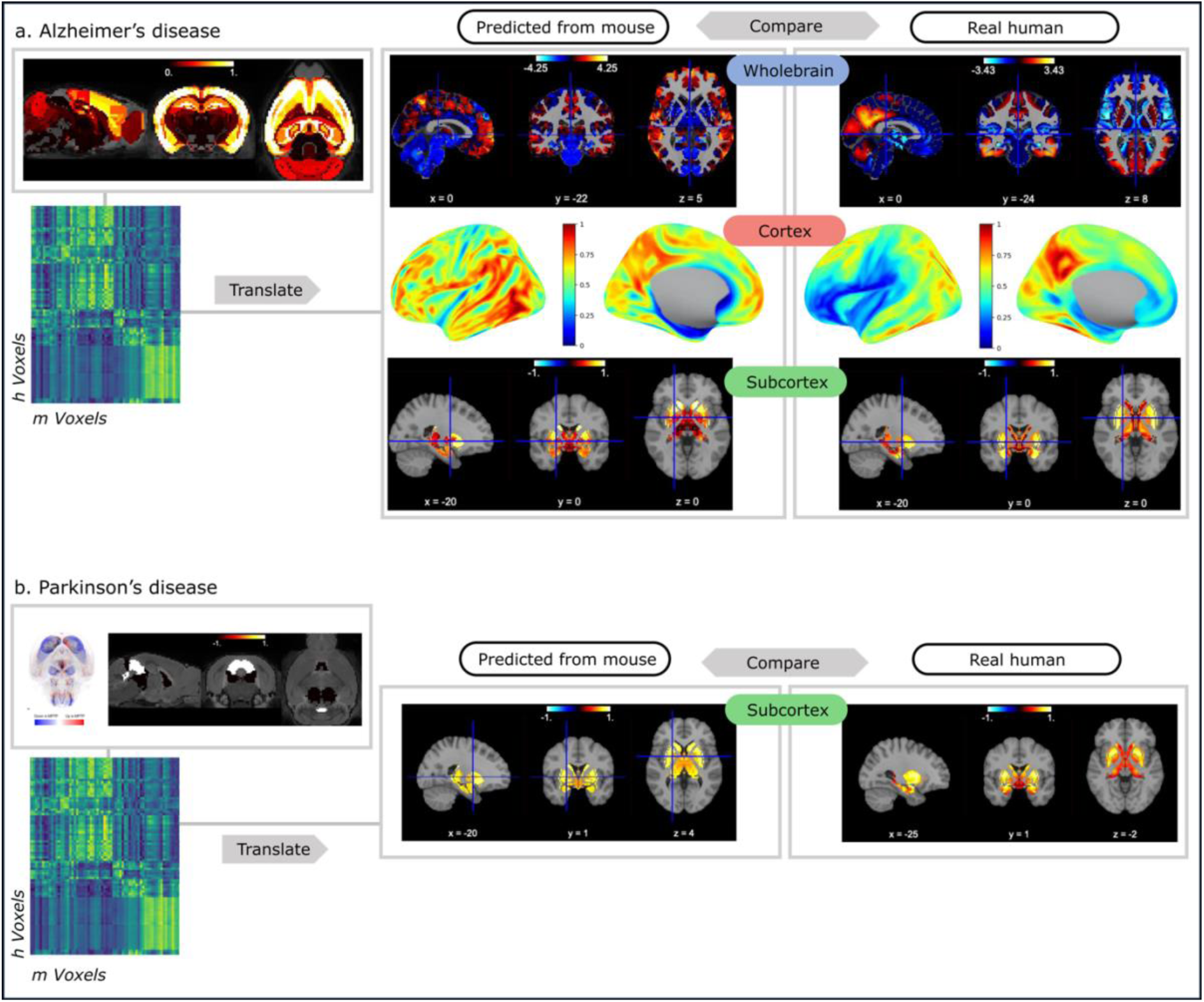
a. Translation of an existing map of plaque deposition in a mouse model of Alzheimer’s disease (APP-PSI, 13 months), divided between whole brain, cortical only, and subcortical only translations. b. Subcortical translation of a mask of regions showing an effect of disease based on an MPTP mouse models of Parkinson’s disease.

We first translated the map of amyloid deposition in the mouse brain focusing on cortical areas only. The correlation between the translated whole-brain map and the real whole-brain human map (r=0.10, p<0.001) was significant, but lower than the correlation when looking at the cortical data only (r=0.22, p<0.001). The subcortical map (containing subcortical areas and hippocampal subfields) correlated significantly with the real subcortical data (r=0.18, p=0.008).

From the whole brain comparison plots in Fig 5a, we can see consistencies with the pattern of brain atrophy in the real data in the posterior cingulate cortex, the precuneus and occipital lobe, as well as in the temporal lobe. The main differences can be seen in the cerebellum and in the frontal areas, where the translation remains consistent with the original mouse map showing respectively low and high plaque deposit. The cortical surface representation further emphasises a pattern of lower atrophy in primary areas, both in the real human map and in the translated map, in addition to the high atrophy component in the posterior cingulate cortex. The subcortical plots highlight that despite not reaching significance, the pattern of cortical atrophy is well mirrored in parts of the caudate nucleus and putamen, as well as in the hippocampus highlighting a consistent gradient along the longitudinal axis, with increased atrophy in the head compared to the body and tail.

#### Translating a map of Parkinson’s disease

Quantitative whole brain maps of a disease are not always readily available. Here, we wanted to give a different example of translation, where we leverage less information but still obtain a basis of comparison with an existing human quantitative map. We created a map consisting of subcortical regions where effects of disease were present, from a whole brain 3D light-sheet fluorenscence microscopy image quantifying tyroxine hydroxylase-positive neurons in MPTP mice compared to controls [32]. In this widely used mouse model of Parkinson’s disease, diseased mice exhibited loss of TH signal intensity in the substantia nigra, caudate nucleus, putamen, globus pallidus and subthalamic nucleus, and increased TH signal intensity mainly in the amygdala and hypothalamus. To recapitulate these results, we gave regions with loss of signal a coefficient of - 1, and regions with increased signal a coefficient of +1 and translated this binary summary image.

We compared the translation to a statistical map of PD > Controls (Neurovault.collection:2694,[33]). In this study, functional connectivity differences with seeds in the basal ganglia were investigated between Parkinson’s disease patients and healthy aging controls. Reduced functional connectivity with the basal ganglia network was found and this was improved through medication, highlighting the potential of the method to serve as a relevant biomarker for the disease. In our case, this statistical map provides crucial patterns of a metric with patterns of both increase and decrease due to disease, which is consistent with the translation we performed from the mouse.

Given that the mouse map consisted of subcortical regions, we performed a subcortical translation only, meaning that we took the sub-matrix of the similarity matrix corresponding to the subcortical voxels in both species. We found a significant correlation between the translated subcortical image and the real human data masked to keep subcortical areas only (r=0.38, p=0.039).

### Longitudinal translations determine the best mouse model match for a given patient population

Having established the feasibility of translating whole-brain maps of both AD and PD from the mouse to the human, we next tested whether it is possible to dissociate across different models of the same disease. Therefore, we translated data from three distinct mouse models of AD, each which was assessed at multiple ages of the mice.

The translation across timepoints captures the progression of the disease as described for all three models, APP-PS1, hAPP-J20, and Tg2576 (Fig 6 (a)). Consistent with the levels of plaque deposition in the mouse, the APP-PS1 translation has the highest intensity and the widest spatial spread at 19 months. The translation also captures the ‘cortex-first’ pattern of disease progression in the APP model, followed by a spread to the subcortex and cerebellum. The translations also capture the ‘subcortical-first’ pattern of disease progression in the other two models. Particularly in the hAPP-J20 translations, the intensity rises more sharply in the hippocampus, followed by the rest of the brain. The translation of Tg2576 captures the initial slow progression of the disease until 9 months, followed by an increase that overtakes the progression of the hAPP model.

**Fig. 6.**
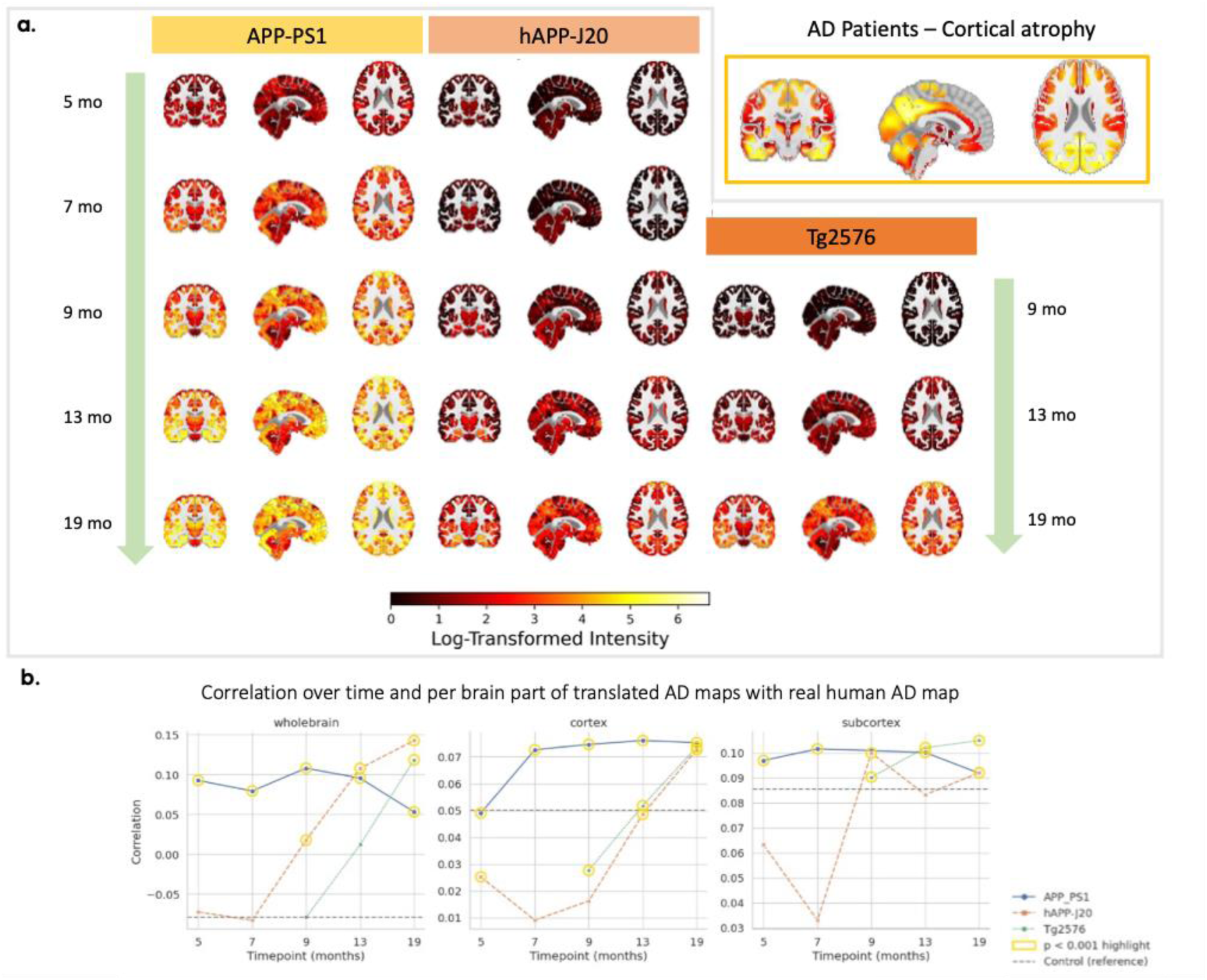
a. Translation of plaque deposition in three mouse model of AD (APP-PSI, hAPP-J20, and Tg2576) at multiple timepoints, compared with human patient data of cortical atrophy in AD. b. Correlation scores across timepoints between each translated mouse model and human cortical atrophy in AD, across different brain masks.

Looking on average at the correlations with the human patient cortical atrophy, the APP model starts off as the best match for the human data, but this changes at 13 months, as the disease progresses in the other two models. The whole brain correlation between APP-PS1 and human AD decreases after 13 months, the mice potentially having reached a more severe disease stage than the humans, described as having ‘mild dementia’.

Taken together, there is a trend that is captured by the translation, for all mouse models to become increasingly correlated with the human data as the disease progresses especially in the cortex. For whole-brain effects, our results suggest that after 13 months hAPP-J20 may be the best reflection of the human phenotype, while Tg2576 may better reflect the subcortical phenotype.

## Discussion

Translational neuroscience suffers from a lack of formal comparative models. To interface between the mouse and the human, separated by over 85 million years of evolution, we built a common space based on the spatial patterns of expression of gene orthologs. Using this common space, we first demonstrated the ability of our approach to identify known cross-species homologies, before investigating how mouse regions translate across the much-expanded human neocortex, highlighting a one-to-one mapping of regions in all major anatomical subdivisions, as well as broader yet anatomically meaningful matches across regions within these subdivisions. Moving beyond anatomical comparisons, we exploited the common space to predict patterns of cortical atrophy in Alzheimer’s disease based on a map of plaque deposition in the mouse brain [31], and we also found a significant correlation between the translation of MPTP mouse map of disease effects and differences in functional connectivity in the human. Finally, we determined the best match for a given human phenotype across three widely used mouse models of AD and multiple timepoints per model.

Our common space took advantage of the availability of whole-brain transcriptomic maps in both the human and the mouse [8], [9]. Patterns of spatial gene expression have previously been used to show similarities in brain organization across the mouse and the human [39] and compare the effects of sex differences across the species [40]. The present work formalized this approach by in effect describing both the mouse and human brain in a common transcriptomic space, similar to what has previously been done for primate brains in connectivity space [15], [41]. However, the strong autocorrelation across genes and the sheer size number of data features mean that a data reduction is likely to improve the common space [19], [42]. We used such an approach in our earlier work [19], by training a multi-layer perceptron (MLP), and providing the foundation for quantitative, whole brain comparison of the mouse and human brain based on cross-species embeddings. However, the MLP did not by design enforce cross-species alignment: the cross-species similarity lacked specificity across broad anatomical areas such as the cortex and the cerebellum.

The present study uses a variational autoencoder to leverage the specific characteristics of its latent space. We trained the model on the mouse data, which is both the simplest of the two brains—preventing overfitting—and the ‘model’ species based on which inference need to be made in translational neuroscience. Pragmatically, the coverage of the mouse transcriptomic data was also complete and high resolution, while in the human dataset this was achieved through interpolation. In our approach, the VAE itself provided us with a handle on the level of clustering versus scattering of the latent vectors. By combining the VAE with a latent classifier, we forced the latent space to remain sensitive to regional boundaries. We were thus able to achieve a balance between cross-species alignment and within-species regional clustering, in a semi data-driven way. As a result, our approach achieved significant improvement of the mouse human comparison compared to earlier approaches. We note that an alternative approach was taken by a recent study [20], but this required incorporation of additional receptor data as well as structural priors in a graph-based architecture.

We provide the first systematic translation of binary regions across the entire mouse brain, yielding human voxel-wise distributions of values of similarity to each mouse region. We highlight the successful correspondence of most known homologs, with focal matching across all major anatomical areas. Standing out among the most focal translations, our hippocampal results are consistent with recent studies of cross species alignment [42], and in depth comparison of macaque and human hippocampus [43]. This translation highlights increased similarity in the head of the hippocampus, but across all subfields, which is consistent with patterns of molecular variation along the longitudinal axis of the hippocampus, rather than the distal-proximal, as documented in the human (Vogel et al., 2020).

We also highlight a number of one-to-many correspondences, in the cases where the translation spans beyond its supposed homologous area, with meaningful underlying patterns of brain organisation. Conserved brain organisation principles are a strong advocate for the successful use of mouse models, even if the areas in the human brain have appeared after the separation of rodent and primate lineages, as was demonstrated in a recent study comparing the mouse and human premotor cortex [44]. In our case, the beyond-target translations always remain within the main anatomical constituent tissue, and are mainly found in the cortex. They reveal two complementary patterns of brain organisation when looked at in combination. The first is a gradient between primary and association cortices, which is a well-known finding, replicated across modalities and species [36], [39], [45]. The second is a modality-specific pattern whereby primary sensory areas have higher similarity to sensory association areas corresponding to the same modality.

Ultimately, the goal of our approach is to establish a quantitative footing for translational neuroscience, by enabling maps of brain organization or brain changes due to disease or treatment to be directly compared across species by translating through the common space. Here, we focused on two examples of neurodegenerative diseases, where translation rates from the preclinical setting to patient population are particularly low, and for which the data is heterogeneous across species. We demonstrate our model’s ability to translate statistical results originating from different data modalities.

In Parkinson’s disease, the changes observed in the brainstem nuclei, with the loss of dopaminergic neurons in the substantia nigra, lead to compensation mechanisms in the basal ganglia and to the disruption of its networks [33]. We were able to retrieve this result by translating a binarized map of statistical differences reported between MPTP mice and wild type mice, even in the absence of a full, continuous, quantitative statistical map. The translation of disease effects in the mouse correlated significantly with changes in resting state functional connectivity in Parkinson’s disease patients. This result holds promise in terms of being able to make inferences even when the best possible data is not readily available for translation.

In humans, one aspect of Alzheimer’s disease is amyloid plaque deposition, which has been shown to start in the cortex [46], progressing to the hippocampus, striatum, basal forebrain, thalamus, and to arise at the later stages in brainstem nuclei and cerebellum [47], [48], [49]. Based on this spatial progression profile, the APP-PS1 model has been suggested to best mirror the later stages of the disease [31], which is why we started by translating this model. As the patient population that we were comparing to were described as presenting ‘mild dementia’, we chose an intermediate timepoint – 13 months. We found that it correlated significantly with human cortical and subcortical atrophy maps [38]. This result prompted us to extend the approach to simultaneous comparison of three mouse AD models, at multiple timepoints. We confirm that the APP-PS1 mouse model is likely better suited for advanced stages of the disease, as the spatial spread is extensive and amounts of plaque deposition are largely predominant over the other two models. However, after 13 months, the difference between the models is less clear cut, with hAPP-J20 potentially better matching a ‘mild dementia’ patient population, and Tg2576 better representing patients with later but more rapid spread, and with extensive subcortical disease development.

The increasing availability of neuroimaging-derived phenotypes in both mice [50], [51] and humans [52], [53] entails that the data are now available to scale our approach toward multi-level comparisons. While matching data modalities is often sought after in comparative studies, here we prove that we can manage heterogeneity and thus harness state of the art approaches for each species. Increasing data from mouse models and human disease will allow more direct comparisons, including elucidating where models fail and which aspects of a disease are captured by the variety of available models [54]. Potentially more impactful even is the ability to link groups of patients in heterogenous syndromes, such as often seen in neurodevelopment and psychiatry, where clustering approach have been able to identify subtypes both in humans and mice [55]. In these cases, mouse models of a disease often only capture a particular variant or aspect of its etiology; direct between-species comparisons have the potential to link models to specific (sub)groups of human patients.

In conclusion, we highlight a highly flexible and scalable framework to support translational neuroscience investigations ranging from fundamental enquiries to disease outcome prediction. Preclinical research is leaning towards more diverse animal models, while interest in personalized medicine is growing in the clinical setting.

We believe that our approach takes a clear step towards answering the question: ‘what is the best animal model for this patient?’

## Author contributions

*Conceptualization: CJ, SNS, RBM*

*Methodology: CJ, MA, AB, SNS, RBM*

*Funding acquisition: RBM, JPL, SNS*

*Supervision: RBM, JPL, SNS*

*Writing – original draft: CJ, RBM*

*Writing – review & editing: CJ, MA, AB, JPL, SNS, RBM*

